# Characterization of 100 sequential SARS-CoV-2 convalescent plasma donations

**DOI:** 10.1101/2020.06.21.163444

**Authors:** Christof Jungbauer, Lukas Weseslindtner, Lisa Weidner, Simon Gänsdorfer, Maria R. Farcet, Eva Gschaider-Reichhart, Thomas R. Kreil

**Affiliations:** Austrian Red Cross, Blood Service for Vienna, Lower Austria and Burgenland, Vienna, Austria; Center for Virology, Medical University of Vienna, Vienna, Austria; Global Pathogen Safety, Baxter AG, a Takeda Company, Vienna, Austria

## Abstract

Transfusion of SARS-CoV-2 convalescent plasma is a promising treatment for severe COVID-19 cases, with success of the intervention based on neutralizing antibody content. Measurement by serological correlates without biocontainment needs, and an understanding of donor characteristics that may allow for targeting of more potent donors would greatly facilitate effective collection.

## Text

The transfusion of convalescent plasma (CP) for the treatment of infectious diseases has been in medical use for more than 100 years and was found worthy of the first Nobel Prize in Medicine, awarded to Emil von Behring in 1901. Based on some earlier successes of CP therapy with the other two zoonotic coronaviruses that have caused large numbers of severe respiratory infections in humans during the last two decades, i.e. SARS-CoV and MERS-CoV (1, 2), its use was quickly initiated after the emergence of the now pandemic SARS-CoV-2 in late 2019 (3-5).

For the identification of potential SARS-CoV-2 CP donors, different analytical approaches are possible, and some have been suggested (6, 7). These include detection of the virus by nucleic acid testing during acute infection, and after convalescence the detection of virus binding antibodies by ELISA, lateral flow or Western blot assays, or the detection of functional antibodies by virus microneutralization test (MNT)(8). Only a MNT provides for a functional correlate of antiviral efficacy and this assay is therefore still considered as ‘gold standard’ in serological testing, yet testing needs to be conducted under level 3 biosafety containment restrictions (BSL3), requires a few days to generate results and is more complex to perform. It would thus greatly facilitate testing for SARS-CoV-2 antibodies in potential plasma donors and their donations, if a more simple and easily scalable binding assay can be correlated against functional antibody activity as determined by MNT.

In addition, CP collections might become more effective if donor characteristics were understood to correlate with higher functional antibody activity, such as e.g. disease severity, days between disease onset and plasma collection, or age and sex of donors.

The Austrian Red Cross Blood Service for Vienna, Lower Austria and Burgenland initiated the collection of virus-inactivated CP by plasmapheresis at the Vienna Blood Centre and the first 100 CP units collected were tested for functionally active neutralizing antibodies by MNT (Attachment). While the absolute numbers generated by any such assay need to await availability of international reference standards before any meaningful comparison becomes possible, on average the CP units had high neutralizing antibody titer, with a mean microneutralization titer 50% (NT_50)_ of approx. 1:230 (Figure 1A – NT_50_ histogram, median 1:100, 25% percentile 1:54, 75% percentile 1:231, range < 1:8 – 1:1765). Somewhat surprisingly for individuals who had successfully recovered from NAT-confirmed SARS-CoV-2 infection, neutralizing antibodies were undetectable in one plasma unit, and for a total of six units (6%) the NT_50_s were below 1:23, i.e. INV log_2_ NT_50_ 4.5 (Figure 1A).

**Figure 1.**
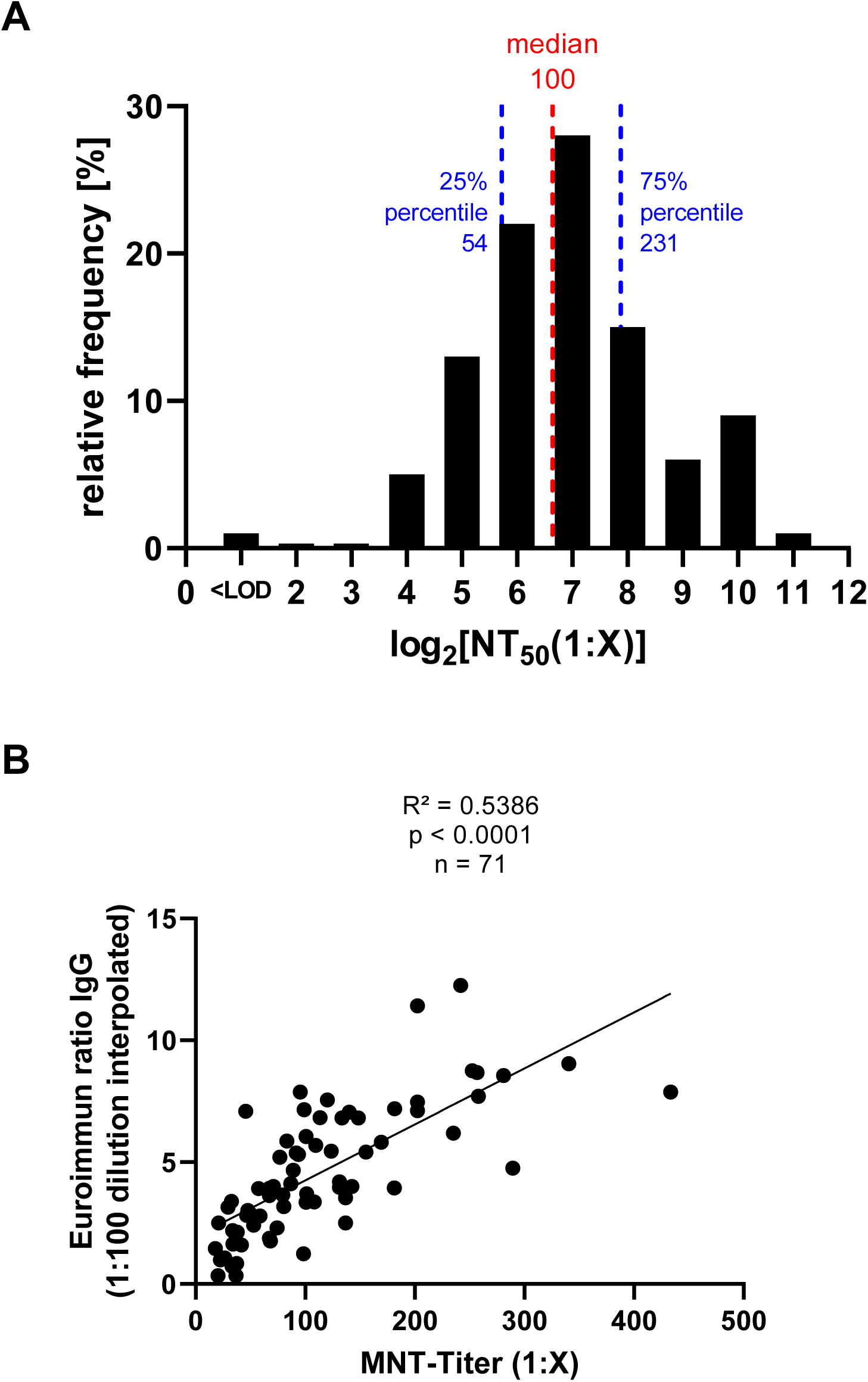
Characterization of 100 convalescent plasma donations collected at the Vienna Blood Centre for (**1A**) SARS-CoV-2 neutralizing antibody content, reported as log_2_ microneutralization titers 50% (1:X) plotted against the relative frequency (%) of occurrence. (**2A**) NT_50_ titers were correlated against Euroimmun IgG ELISA signal ratios obtained for the same donations.

A binding antibody assay was also performed on these CP units (Attachment), with the intention to establish a correlation between the functionally more relevant MNT and the more accessible ELISA, as well as an ELISA threshold to allow for the elimination of sub-potent units and the qualification of CP units for transfusion. For the totality of the samples for which valid results from both assays were available (N = 83), the correlation between results from the MNT and the ELISA was highly significant (p < 0.0001), yet quantitatively limited (R^2^ = 0.2830) (Figure 1B). With the intention of using the ELISA primarily for establishing a lower threshold for CP units to issue them for treatment of COVID-19 cases, the analysis was repeated excluding particularly high NT_50_ titers (>1:500, N = 12 / 14.5%), which improved the correlation (R^2^ = 0.5386; Figure 1B). Using the Euroimmun ELISA 1.1 cut-off for positivity to qualify units for transfusion, 6 units (7.2%) with an average NT_50_ of 1:29 would have been excluded, with a corresponding increase of the mean NT_50_ of all collected plasma units from 1:233 (N = 83) to 1:249 (N = 77). A final verdict about whether this cut-off is suitable for use of CP in the treatment of COVID-19 will have to await an evaluation of clinical efficacy in correlation to these antibody measurements.

For the targeted collection of high antibody titer CP units it would be helpful to understand donor characteristics that might correlate with higher antibody potency, for example it has been suggested that increasing disease severity may result in the development of higher antibody titers (9). Of the 100 plasma donors, 90 were classified into WHO disease severity scores of 1 and 2 (10), with an average NT_50_ of 1:208, versus only 6 donors in disease severity scores of 3 to 6, who had a mean NT_50_ antibody titer of 1:696. While these results may seem to support the notion of higher titers with increased disease severity, the numbers of CP donors in higher WHO scores in this study are too low to determine significance. For CP donor age (Figure 2A), the NT_50_s were significantly correlated (p = 0.0019), yet with little predictive value (R^2^ = 0.09466) and targeting advance age donors for CP collection may therefore not be very effective. On average, the CP collected from male donors (N = 61) had a significantly higher mean NT_50_ than from female donors (N=38), yet whether the mean titer difference of 1:220 has any functional relevance is questionable (Figure 2B).

**Figure 2.**
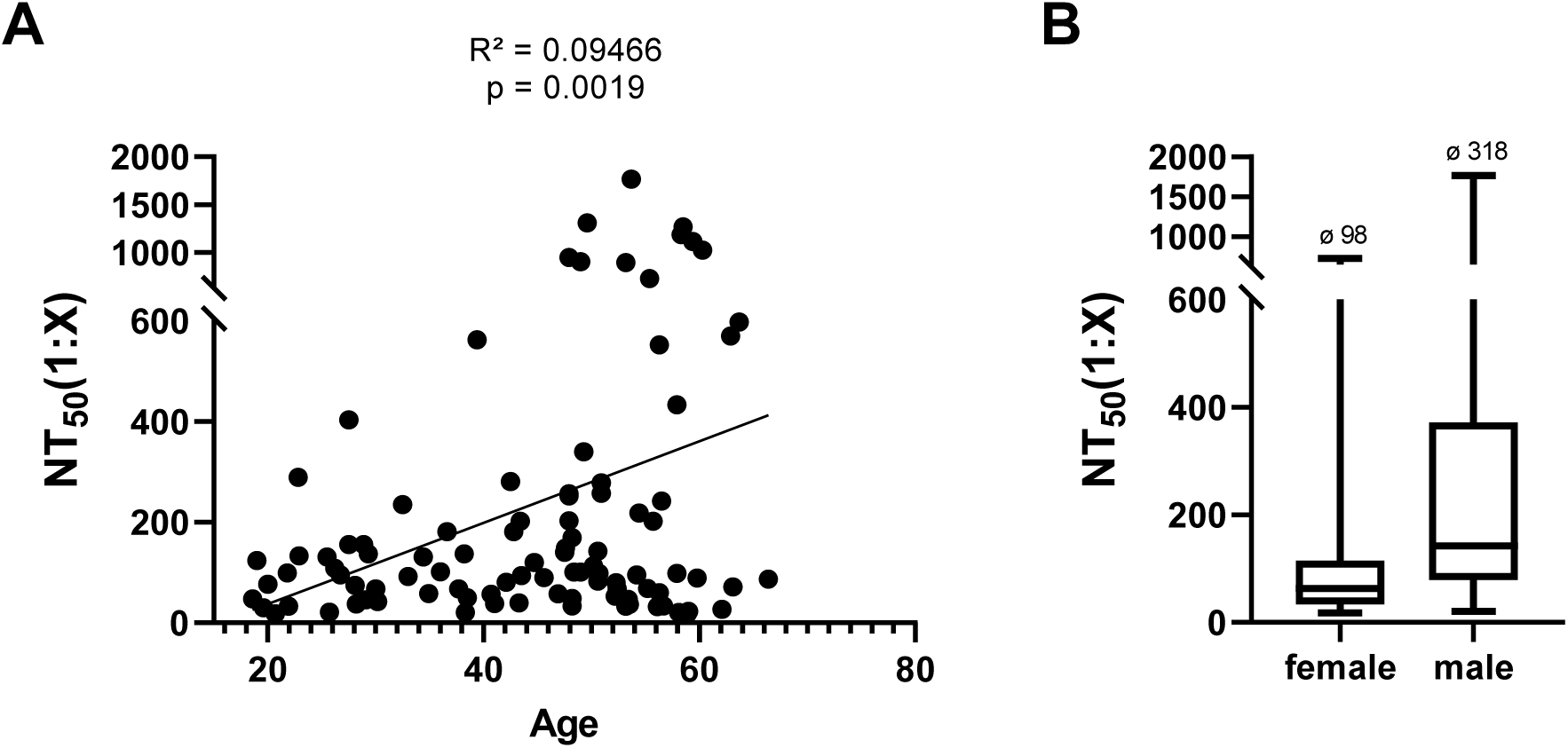
COVID-19 convalescent plasma donor demographics illustrating (**2A**) correlation of SARS-CoV-2 neutralizing antibody titers (NT_50_) with donor age and (**2B**) difference in NT_50_ between female and male donors.

## Conclusions

While functional testing for SARS-CoV-2 neutralizing antibodies would be desirable for CP units before issuing them for treatment of COVID-19, biosafety requirements and considerations around assay duration and complexity make this approach impractical. An adequate pooling strategy of convalescent plasma may level out variations of antibody titers and quality in the therapeutic units. Here we have established an ELISA-based correlate to the MNT, with a threshold proposal that could be used to eliminate lower titer units from the clinical supply for COVID-19 treatment. Disease severity may be associated with the development of higher titers of neutralizing antibodies, although larger case numbers will be needed for a final conclusion. And while age and gender are significantly correlated with MNT antibody levels, the differences are small and thus probably not helpful for CP donor targeting.

## Acknowledgments

The contributions of the Global Pathogen Safety team from Takeda, most notably Melanie Graf, Simone Knotzer, Alexandra Schlapschy-Danzinger, Eva Ha, Elisabeth List, Nicole Rameder, Michael Karbiener, Petra Gruber, Stefan Pantic, Michaela Rumpold and Julius Segui (neutralization testing) as well as Veronika Sulzer and Sabrina Brandtner (cell culture) are gratefully acknowledged. Andrea Hörl and Hannah Griebler (Center for Virology, Medical University of Vienna) are gratefully acknowledged for ELISA testing. SARS-CoV-2 was sourced via EVAg (supported by the European Community) and kindly provided by Christian Drosten and Victor Corman (Charité Universitätsmedizin, Institute of Virology, Berlin, Germany).

## Disclaimers

MRF, EG-R and TRK are employees of Baxter AG, Vienna, Austria, now part of the Takeda group of companies. MRF and TRK have Takeda stock interest.

## Author Bio

(first author only, unless there are only 2 authors)

Dr. Jungbauer is medical doctor and specialist in immunohematology and transfusion medicine. He currently is medical director of the Austrian Red Cross Blood Service for the East Region, based in Vienna, Austria. His research interests are virus diseases and molecular cell and tissue antigen typing.

