## Supplementary Material and Methods for "Characterization of 100 sequential SARS-CoV-2 convalescent plasma donations"

### Materials and Methods

#### Collection and virus-inactivation of CP

All CP donors had PCR-confirmed SARS-CoV-2 infections. Plasma was collected at least 28 days after the end of clinical symptoms in concordance with all applicable legal requirements and relevant recommendations. The donors signed an informed consent regarding CP and special testing.

The plasma was collected using Trima Accel® apheresis version 7.0 devices (Terumo, Eschborn, Germany) with Trima Accel® MultiPlasma sets (ref. 82700). The collection volumes ranged from 440 to 650 ml adapted to bodyweight of the donors. Samples of each donation were investigated for SARS CoV-2 antibody titers by neutralization assay and Euroimmun Anti-SARS-CoV-2-ELISA (IgG) (Euroimmun, Lübeck, Germany).

CP was pooled (at least two plasmas per pool) based on the neutralization assay titer results yielding for final antibody titers of >= 1:300 in the therapeutic units. Subsequently the pooled plasma was pathogen inactivated using Intercept® Processing Sets for Plasma (Ref. INT3104B-1, Cerus Europe, Amersfoort, Netherlands) with the Intercept® UVA-Illuminator (Ref. INT100). The final volume of the therapeutic CP units was adjusted to 200 ml. At time of submission of this manuscript twelve patients had received 27 units (ranging from 1 to 3 units per patient, median 2.0 units) in Vienna and Lower Austria.

### SARS-CoV-2 infectivity assay and testing of CP for neutralizing antibodies

SARS-CoV-2 strain BetaCoV/Germany/BavPat1/2020 was kindly provided by the Charité Universitätsmedizin, Institute of Virology, Berlin, Germany; EVAg 026V-03883. Vero cells (ATCC CCL-81), sourced from the ECACC (84113001) were cultured in TC-Vero medium supplemented with 5% FCS, L-glutamine (2mM), nonessential amino acids (1x), sodium pyruvate (1mM), Gentamycin sulfate (100 mg/ml) and sodium bicarbonate (7.5%). For determination of SARS-CoV-2 infectivity by tissue culture infectious dose 50% ($TCID_{50}$) assay, eightfold replicates of serial half-log sample dilutions were incubated on cells for 5-7 days before any cytopathic effect (CPE) was assessed microscopically. Virus concentrations were calculated according to the Poisson distribution.

For virus microneutralization assays, CP samples were serially 1:2 diluted and incubated with 100 $TCID_{50}$ of SARS-CoV-2. The samples were subsequently applied onto Vero cells seeded in tissue culture microplates and incubated for 5-7 days, when cells were evaluated for the presence of a CPE and the SARS-CoV-2 microneutralization titer ($NT_{50}$), i.e. the reciprocal sample dilution resulting in 50% virus neutralization, was determined using the Spearman-Kärber formula.

### Testing of CP for binding antibodies by ELISA

Anti-SARs-CoV-2-IgG antibodies were quantified by the Euroimmun SARS-CoV-2 IgG ELISA (Euroimmun, Lübeck, Germany) using an Euroimmun Analyzer I with the protocol recommended by the manufacturer. IgG concentration is given as Ratio, calculated using optical density and an internal calibrator provided by the manufacturer. The Euroimmun SARS-CoV-2-IgG ELISA uses the recombinant structural protein (S1 domain) of the spike protein as antigen. Serum samples were tested in serial dilutions of twofold steps from 1:5 to 1:1280. When the IgG Ratio in a sample still exceeded 1.1 (the cut-off recommended by the manufacturer) at the dilution of 1:1280, further dilutions (1:5120 to 1: 20480) were performed for an accurate interpolation.
